# Altered systemic bioenergetic reserve in chronic kidney disease predisposes hearts to worse functional outcomes

**DOI:** 10.1101/2024.10.24.620055

**Authors:** Megan Young, Malene Aastrup, Nikayla Patel, Fenn Cullen, Esben S.S. Hansen, James E. Clark, Thomas R. Eykyn, Michael Vaeggemose, Ana Vujic, Loucia Karatzia, Ladislav Valkovič, Jack J.J.J. Miller, Niels H. Buus, Christoffer Laustsen, Magdi M. Yaqoob, Dunja Aksentijevic

## Abstract

**Background:** Cardiovascular mortality in chronic kidney disease (CKD) remains disproportionately high, yet the mechanisms linking renal dysfunction to cardiac vulnerability are incompletely understood. Uraemic cardiomyopathy is increasingly recognised as a systemic metabolic disease, but the contribution of multi-organ bioenergetic failure in cardiac dysfunction is poorly defined.

**Hypothesis:** CKD induces metabolic remodelling across peripheral organs (liver, skeletal muscle, and kidneys) depleting systemic bioenergetic reserve, compromising cardiometabolic flexibility and stress resilience.

**Methods:** Using CKD models of different aetiologies in rats (glomerulosclerosis by partial nephrectomy and interstitial fibrosis by adenine diet) we investigated cardiac and systemic metabolic remodelling.

**Results:** Irrespective of aetiology, renal insufficiency resulted in cardiac dysfunction including impaired functional recovery after 25-minutes ischaemia. ^1^H NMR metabolomic analysis revealed perturbations of systemic metabolism in CKD were more severe than cardiometabolic changes with alterations of skeletal muscle, liver, and kidney metabolism indicating reduced systemic bioenergetic reserve. This pre-clinical observation was recapitulated in human CKD patients where phosphorus magnetic resonance spectroscopy assessment of exercising lower leg muscle identified bioenergetic deficiencies preventing maximal force generation. Thus, both heart and skeletal muscles in CKD have impaired response to metabolic stress.

**Conclusions:** CKD induces multi-organ metabolic failure that limits the hearts ability to meet energetic demands under stress. This study identifies systemic bioenergetic collapse as a contributing factor to uraemic cardiomyopathy, thus targeting peripheral organ metabolism may represent a novel therapeutic strategy to improve cardiac outcomes in CKD.

**Key learning points:** *What Was Known:* CKD significantly increases cardiovascular risk, but the cause of heart failure in these patients is largely attributed to cardiac pathology alone whilst the potential contribution of systemic metabolic dysfunction remains unexplored.

*What This Study Adds:* Utilising clinical and pre-clinical approach we show that CKD triggers widespread metabolic dysfunction in the liver, skeletal muscle, and kidney, depleting the systemic bioenergetic reserve and impairing the heart’s ability to handle metabolic stress.

*Potential Impact:* These findings show uraemic cardiomyopathy is a **multi-organ metabolic disease** and targeting peripheral metabolic dysfunction could offer a **new therapeutic strategy** to enhance cardiac resilience by restoring systemic energy balance.

## INTRODUCTION

Chronic kidney disease (CKD) affects 840 million people globally and is a leading cause of mortality [1]. CKD progression generates a high-risk clinical phenotype, including inflammation, malnutrition, altered autonomic and central nervous system activity, vascular and bone disease [2]. However, for many CKD patients the risk of developing cardiovascular disease is higher than progression to kidney failure [3]. Uraemic cardiomyopathy is a distinct pathology characterised by diastolic dysfunction, left ventricular hypertrophy (LVH), fibrosis and maladaptive changes in cardiac metabolism [4]. In severe CKD, cardiac complications can progress into heart failure (HF), largely with preserved ejection fraction (HFpEF) [4, 5]. Previous CKD research shows that skeletal muscle energetic reserve mirrors cardiac energetic reserve [6], highlighting the interconnecting cross-talk between two physiological systems. Given that the driving factors for uraemic cardiomyopathy are multifactorial, it remains unknown whether cross-talk between systemic metabolic remodelling in CKD could be contributing to uraemic cardiomyopathy.

CKD is an inherently systemic disease, renal insufficiency causes dysfunction of multiple organs [2]. In addition to perturbed kidney metabolism, extensive systemic metabolic alterations accompany CKD including insulin resistance associated with increased fatty acid catabolism and muscle wastage [7]. Skeletal muscle is a crucial organ for metabolic homeostasis, playing an essential role in insulin sensitivity, protein metabolism, and energy expenditure. However, individuals with CKD often experience alterations in muscle metabolism that can exacerbate their clinical condition and contribute to cachexia. Furthermore, reduced mass and strength contribute to fatigue suffered by CKD patients which notably impairs quality of life [8].

Alterations in renal clearance significantly impact liver function triggering altered carbohydrate and bilirubin metabolism as well as hepatic dyslipidemia [9]. The hepatic-renal interorgan cross talk is vital to systemic metabolic homeostasis and detoxification of the products of protein metabolism, including urea and creatinine [10]. As renal function declines, the accumulation of uraemic toxins can overwhelm the liver’s capacity to metabolise and eliminate toxins, leading to chronic liver damage [11]. This contributes to further deterioration of both liver and kidney function, creating a vicious cycle of comorbidity that can exacerbate CKD patient outcomes [11].

The multi-system nature of CKD justifies an integrated assessment of both systemic and cardiovascular alterations in this population. Combining pre-clinical and clinical approaches we aimed to investigate systemic metabolic changes in CKD and the association with uraemic cardiomyopathy pathophysiology. Most pre-clinical studies use a single animal model of CKD thus poorly mimicking the complexity of clinical CKD.

Our study utilises ***two experimental rat models*** of CKD of different aetiology: adenine diet model which produces rapid-onset kidney disease with extensive tubulointerstitial fibrosis, tubular atrophy, crystal formation and marked vessel calcification, and the partial nephrectomy (PN) model of progressive renal failure due to glomerulosclerosis. In both models, we assessed the impact of CKD development on cardiac function, metabolome, gene expression and susceptibility to ischaemia-reperfusion injury. To assess the impact of kidney dysfunction on systemic metabolism we examined the metabolic profile of plasma, kidneys, liver, and skeletal muscle. Bioenergetic capacity of lower leg muscle was assessed in human CKD patients during exercise using phosphorus magnetic resonance spectroscopy (^31^P MRS).

## RESULTS

### CKD phenotype and cardiometabolic remodelling in CKD animal models

#### Adenine diet induced CKD

0.75% adenine diet CKD induction protocol in Wistar rats led to renal dysfunction evident from increased plasma levels of creatinine and urea compared to control diet (Fig.1A, B) as well as anaemia (Fig.1C). Given the adenine CKD model is characterised by the absence of hypertension [12], there was no increase in LVH index (Fig.1D) or increases in LV wall thickness (Table S2). Changes in left ventricular internal diameter in diastole (LVID,d) and end diastolic volume (EDV) (Fig.1E-F) reflect mild ventricular remodelling and compensatory increases in contractility to maintain ejection fraction (Fig.1G). There is no evidence of congestive HF and lung oedema in adenine CKD (Fig.S2A). mRNA expression of BNP (Fig.1H) as well as PCr/ATP ratio (Fig.1I) were comparable between the adenine CKD heart and controls.

**Fig. 1.**
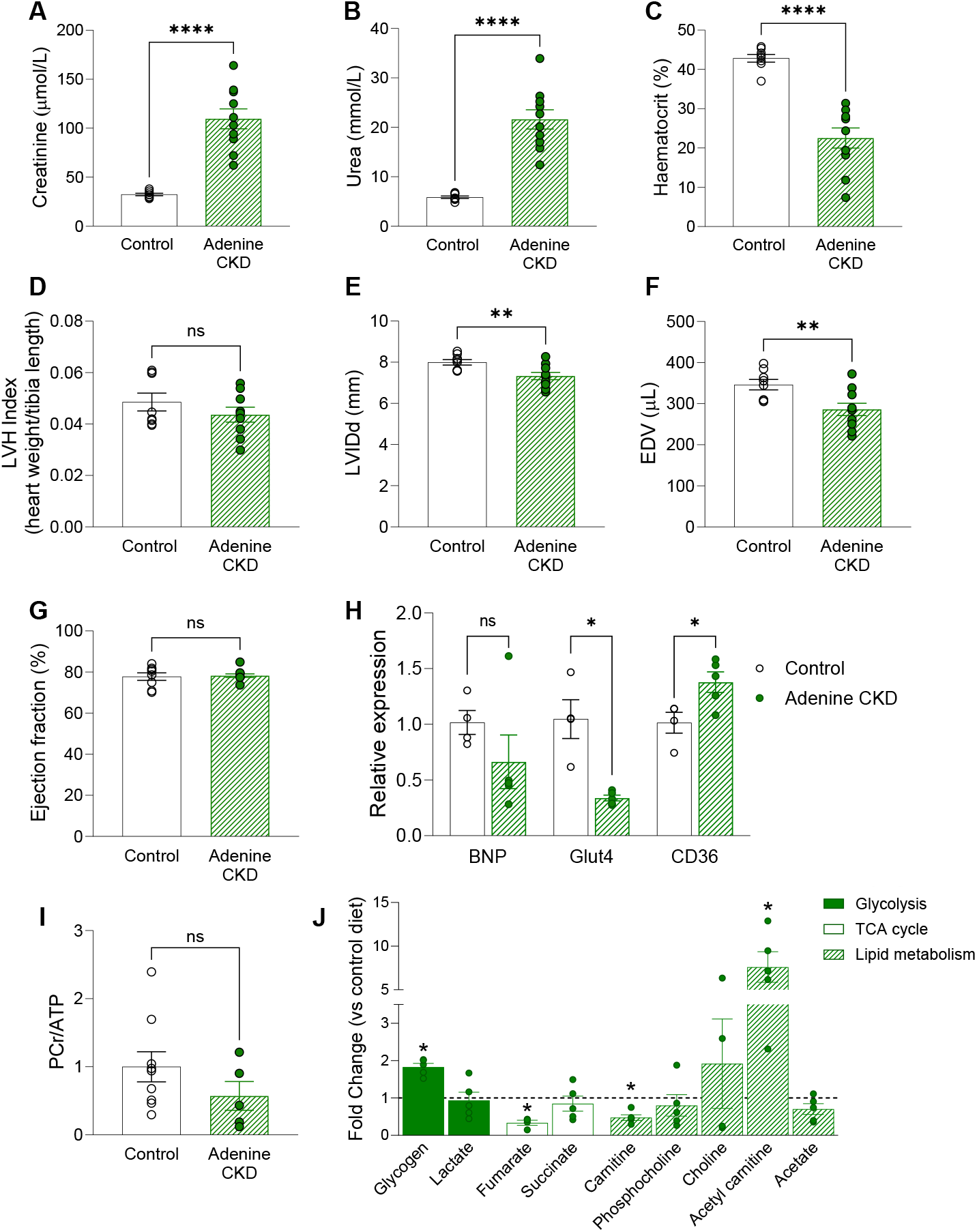
Morphological and cardiac metabolic changes in adenine diet CKD model. (**A-B**) Plasma levels of creatinine and urea (Adenine n=10, control n=8, unequal variance t-test, *p<0.05). (**C**) Haematocrit measurement (Adenine n=10, control n=8, unequal variance t-test, *p<0.05). (**D**) ratio of heart weight to tibia length (Adenine=9, control n=8, Student’s t-test, *p<0.05), (**E-G**) Echocardiography measurement of left ventricle internal diameter in diastole (LVID,d), end diastolic volume (EDV) and ejection fraction (Adenine=10, control n=8, Student’s t-test, *p<0.05). (**H**) mRNA expression of BNP, Glut4 and CD36 relative to housekeeping gene 36B4 (Adenine n=5, control n=4. BNP analysed by Mann-Whitney U test, Glut4 analysed by unequal variance t-test and CD36 analysed by Student’s t-test, *p<0.05). (**I**) Ratio of PCr/ATP measured by ^1^H NMR spectroscopy in cardiac tissue from adenine CKD (n=5) and control diet (n=9) animals. Analysed by Student’s t-test, *p<0.05. (**J**) ^1^H NMR spectroscopy of cardiac tissue from adenine CKD (n=5) expressed as fold change vs control diet (n=9) group. Adenine CKD vs control compared for each metabolite by Student’s t-test, *p<0.05. Data displayed as mean ± SEM.

^1^H NMR spectroscopy assessment of adenine CKD myocardial metabolome revealed perturbed metabolic intermediates related to lipid metabolism, glycogen and TCA cycle (Fig.1J, Fig.S3). Furthermore, adenine diet-induced CKD caused marked changes in cardiac lipid content (Fig.S4). Expression of fatty acid transporter CD36 was increased in adenine CKD hearts, alongside a decreased expression of insulin-dependent glucose transporter GLUT4 (Fig.1H). In terms of plasma metabolic profile (Table 1), adenine diet CKD animals had reduced circulating levels of glucose and insulin compared to control.

**Table 1.** Plasma analysis of adenine diet-CKD and chow diet controls.

|  | Control<br>(n=8) | Adenine CKD<br>(n=10) |
| --- | --- | --- |
| Glucose (mmol/L) | 13.20 ± 0.35 | **10.01 ± 0.36\*** |
| Sodium (mmol/L) | 142.1 ± 0.44 | 140.7 ± 1.17 |
| Potassium (mmol/L) | 4.98 ± 0.21 | **6.58 ± 0.32\*** |
| Chloride (mmol/L) | 95.13 ± 0.69 | 96.40 ± 1.62 |
| Triglycerides (mmol/L) | 1.27 ± 0.28 | 0.82 ± 0.07 |
| Insulin (µg/L) | 1.83 ± 0.24 | **0.54 ± 0.11\*** |
| Free Fatty Acids (µmol/L) | 337.6 ± 62.21 | 224.2 ± 34.01 |
| Alanine aminotransferase (U/L) | 40.14 ± 2.48 | 39.89 ± 2.64 |
| Alkaline Phosphatase (U/L) | 99.00 ± 8.11 | **125.7 ± 8.99\*** |

Analysed by Student's t-test, \*p<0.05 vs. Control. Data displayed as mean ± SEM.

#### Partial nephrectomy (PN) CKD

Two-stage PN surgery induced a CKD phenotype characterised by renal dysfunction (elevated plasma levels of creatinine and urea – Fig.2A-B), anaemia (Fig.2C) and cardiometabolic remodelling. In addition to uraemia and anaemia, hypertension is present in PN model of CKD from the onset [13-15]. 12-weeks of CKD progression led to pathological ventricular remodelling observed from an increase in LVH index (Fig.2D). *In vivo* cardiac assessment identified changes in cardiac wall thickness and decreased internal diameter (Fig.2E-F). Furthermore, elevated mRNA expression of BNP (Fig.2H) is indicative of developing HF however in the absence of a reduction in ejection fraction (Fig.2G). Assessment of circulating metabolites showed a decline in plasma glucose levels in PN CKD (Table 2). ^1^H NMR myocardial metabolomic profiling of PN CKD identified changes in amino acid metabolism (Fig.2J) without alterations in the expression of metabolic transporters (CD36 and Glut4, Fig.2H). However, cardiac energy reserve was impaired as PCr/ATP ratio was significantly reduced (Fig.2I). Comparison of the severity of remodelling in PN vs adenine-diet induced CKD in summarised in Supplementary Figures 5 and 6.

**Fig. 2.**
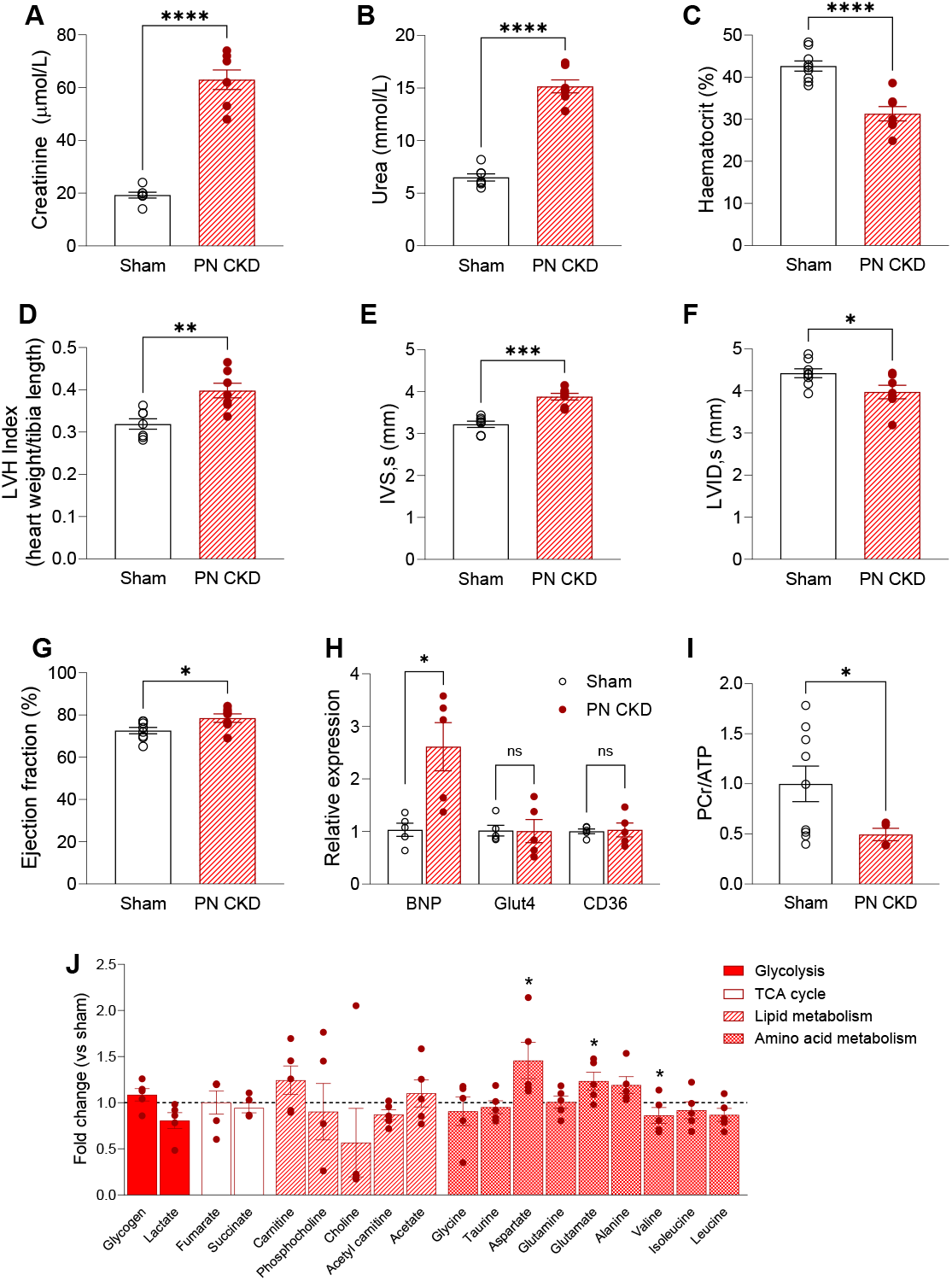
CKD phenotype and cardiometabolic remodelling in PN CKD model. (**A-B**) Plasma levels of creatinine and urea (PN CKD n=7, sham n=7, Student’s t-test, *p<0.05). (**C**) Haematocrit measurement (PN CKD n=7, sham n=9, Student’s t-test, *p<0.05). (**D**) ratio of heart weight to tibia length (PN CKD n=7, sham n=7, Student’s t-test, *p<0.05. (**E-G**) Echocardiography measurement of intraventricular septum thickness in systole (IVS,s), left ventricle internal diameter in systole (LVID,s) and ejection fraction (PN CKD n=7, sham n=8, Student’s t-test, *p<0.05). (**H**) mRNA expression of BNP, Glut4 and CD36 relative to housekeeping gene 36B4 (PN CKD n=5, sham n=5. BNP analysed by unequal variance t-test, Glut4 and CD36 analysed by Student’s *t*-test, *p<0.05). (**I**) Ratio of PCr/ATP measured by ^1^H NMR spectroscopy in cardiac tissue from PN CKD (n=4) and sham (n=9) animals. Analysed by unequal variance t-test, *p<0.05. (**J**) ^1^H NMR spectroscopy of cardiac tissue from PN CKD (n=5) expressed as fold change vs sham group (n=9). PN CKD vs sham compared for each metabolite by Student’s t-test, *p<0.05. Data displayed as mean ± SEM.

**Table 2.** Plasma analysis of PN CKD and sham controls.

|  | Sham<br>(n=7) | PN CKD<br>(n=7) |
| --- | --- | --- |
| Glucose (mmol/L) | 8.3 ± 0.65 | 7.64 ± 0.27 |
| Sodium (mmol/L) | 138.9 ± 1.01 | 138.8 ± 0.4 |
| Potassium (mmol/L) | 4.87 ± 0.18 | 5.58 ± 0.39 |
| Chloride (mmol/L) | 92.43 ± 1.66 | 94.0 ± 0.72 |
| Triglycerides (mmol/L) | 1.5 ± 0.18 | 1.25 ± 0.21 |
| Insulin (µg/L) | 0.79 ± 0.44 | 0.44 ± 0.07 |
| Free Fatty Acids (µmol/L) | 790.1 ± 142 | 757.6 ± 156 |
| Alanine aminotransferase (U/L) | 43.43 ± 3.68 | **32.33 ± 2.41\*** |
| Alkaline Phosphatase (U/L) | 85.57 ± 5.55 | 78.14 ± 7.13 |

Analysed by Student's t-test, \*p<0.05 vs. Sham. Data displayed as mean ± SEM.

### CKD increases susceptibility to cardiac ischaemia reperfusion injury

Given the evidence of altered cardiometabolic profile of CKD hearts, we examined the response to ischaemia and reperfusion. When stressed by a period of 25-minutes total global normothermic ischaemia followed by 25-minutes reperfusion, both adenine and PN CKD hearts display poor outcomes (Fig.3). Given the unaltered PCr/ATP ratio, adenine CKD hearts do show signs of improved recovery during the immediate onset of reperfusion, however this is not sustained (Fig.3). At the end of 25-minutes reperfusion, adenine and PN CKD hearts left ventricle developed pressure (LVDP) recovered 8.02% and 9.75% respectively compared to 50.42% recovery in control hearts (Fig.3).

**Fig. 3.**
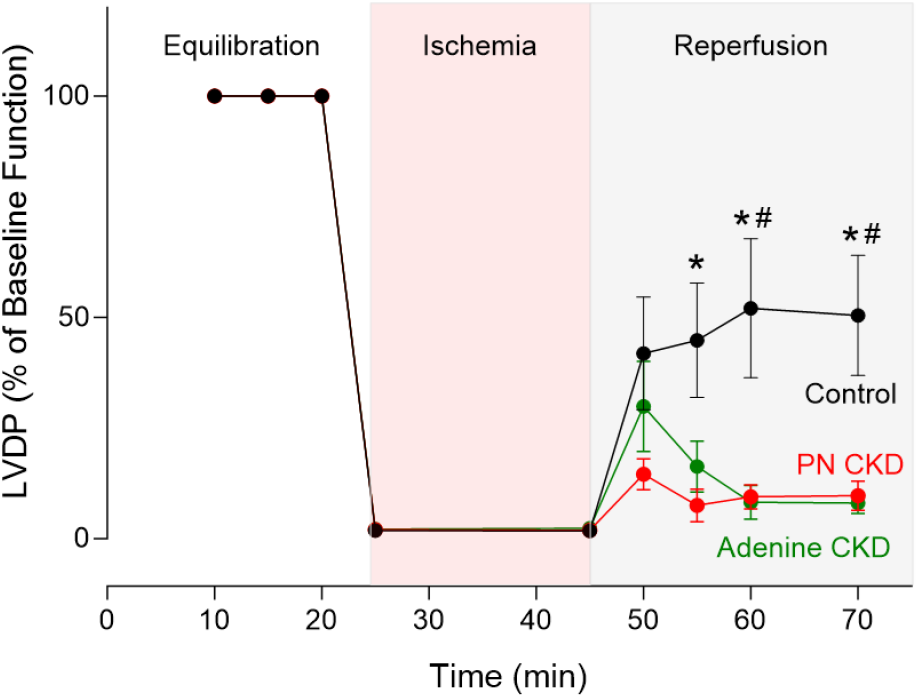
Ischaemia and reperfusion in CKD hearts. Functional recovery in PN CKD (n=4), adenine CKD (n=5) and control hearts (n=5). Left ventricle developed pressure (LVDP) displayed as a percentage of baseline value. Analysed by two-way repeated measures ANOVA, significant interaction (p<0.05) between groups over time. One-way ANOVA analysis for each time point displayed, *p<0.05 PN CKD vs. control, #p<0.05 Adenine CKD vs. control. Data displayed as mean ± SEM.

### CKD development leads to systemic metabolic alterations

#### Skeletal muscle

Skeletal muscle damage was evident in adenine CKD animals and to a lesser extent in PN CKD. Adenine CKD was associated with increased circulating levels of creatine kinase (Fig.4B), as well as a reduction in body weight (Fig.4A) indicative of muscle wastage. The metabolomic profiling of skeletal muscle from adenine CKD animals revealed decline in energy reserve with reduced PCr/ATP accompanied by increased amino acids concentrations (aspartate, alanine, glutamine, glutamate, Fig.4D). The PN CKD model was characterised by doubling of circulating creatine kinase (Fig.4C) and changes in skeletal muscle acetyl carnitine and amino acid profile (Fig.4E).

**Fig. 4.**
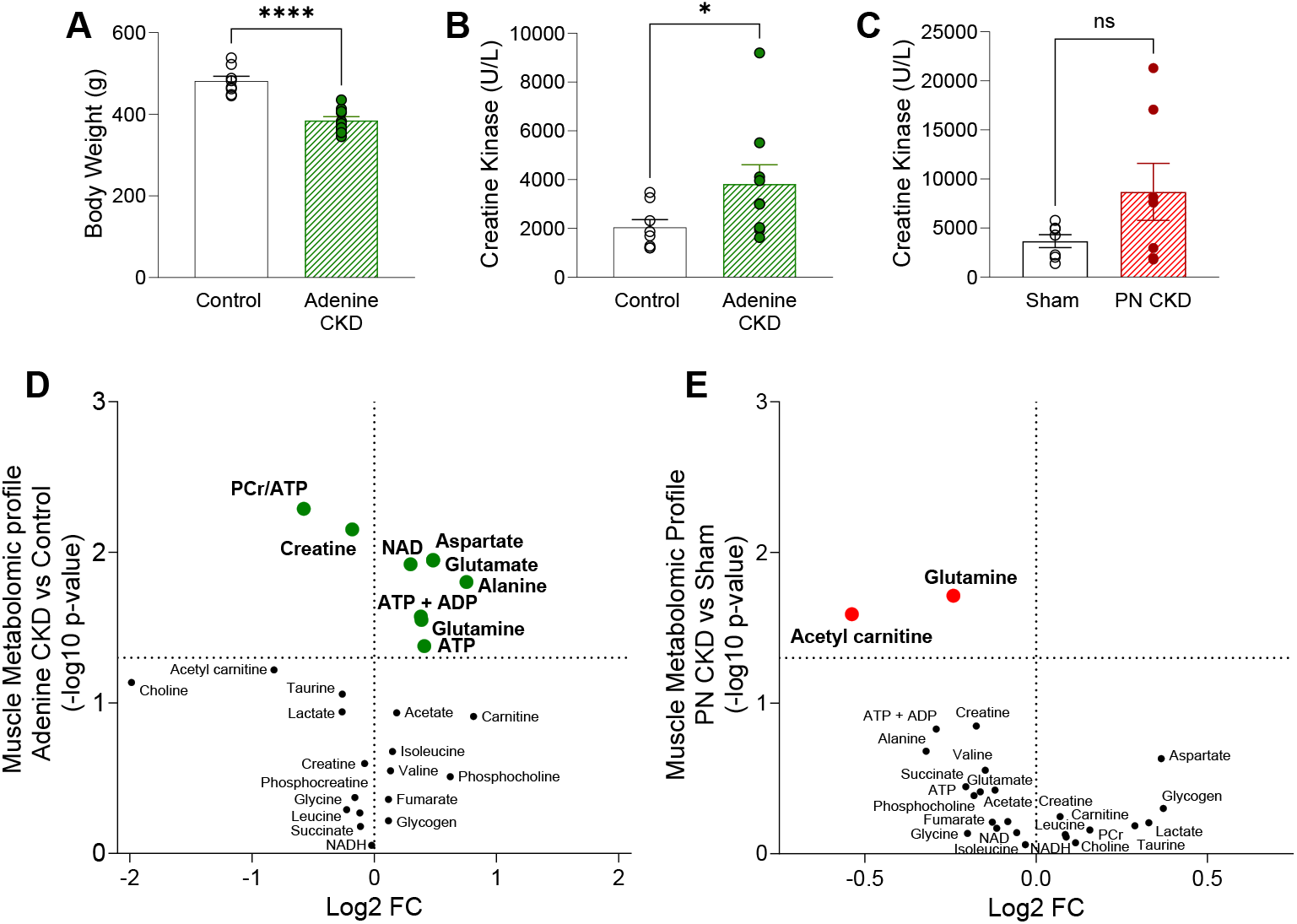
Metabolomic profile of skeletal muscle in adenine and PN CKD models. (**A**) Terminal body weight in adenine CKD animals. Analysed by Student’s t-test, *p<0.05, control n=8, Adenine n=10 (**B-C**) Plasma creatine kinase level in (B) adenine diet CKD animals (Analysed by Mann-Whitney U test, *p<0.05, control n=8, Adenine n=10) and (C) PN CKD animals (Analysed by unequal variance t-test, p>0.05, sham n=8, PN CKD n=8). (**D-E**) Muscle metabolomic profile measured by ^1^H NMR spectroscopy of (D) adenine diet CKD (n=5) vs control diet (n=5) and (E) PN CKD (n=3) vs sham (n=3). Each point represents the fold change (FC) of a metabolite plotted against the associated level of statistical significance for the change analysed by t-test (horizontal dashed line indicates p=0.05 threshold). Data displayed as mean ± SEM.

#### Kidney

Renal dysfunction is evident in both CKD models with increased plasma levels of creatinine and urea (Fig.1A-B, 2A-B). In the adenine CKD model both kidneys are intact accompanied by significant kidney hypertrophy (Fig.5A-B). Kidney morphology is harder to compare with weights in the PN CKD model given the surgical resection, although there is visual evidence of compensatory hypertrophy in the 1/3 remaining kidney.

**Fig. 5.**
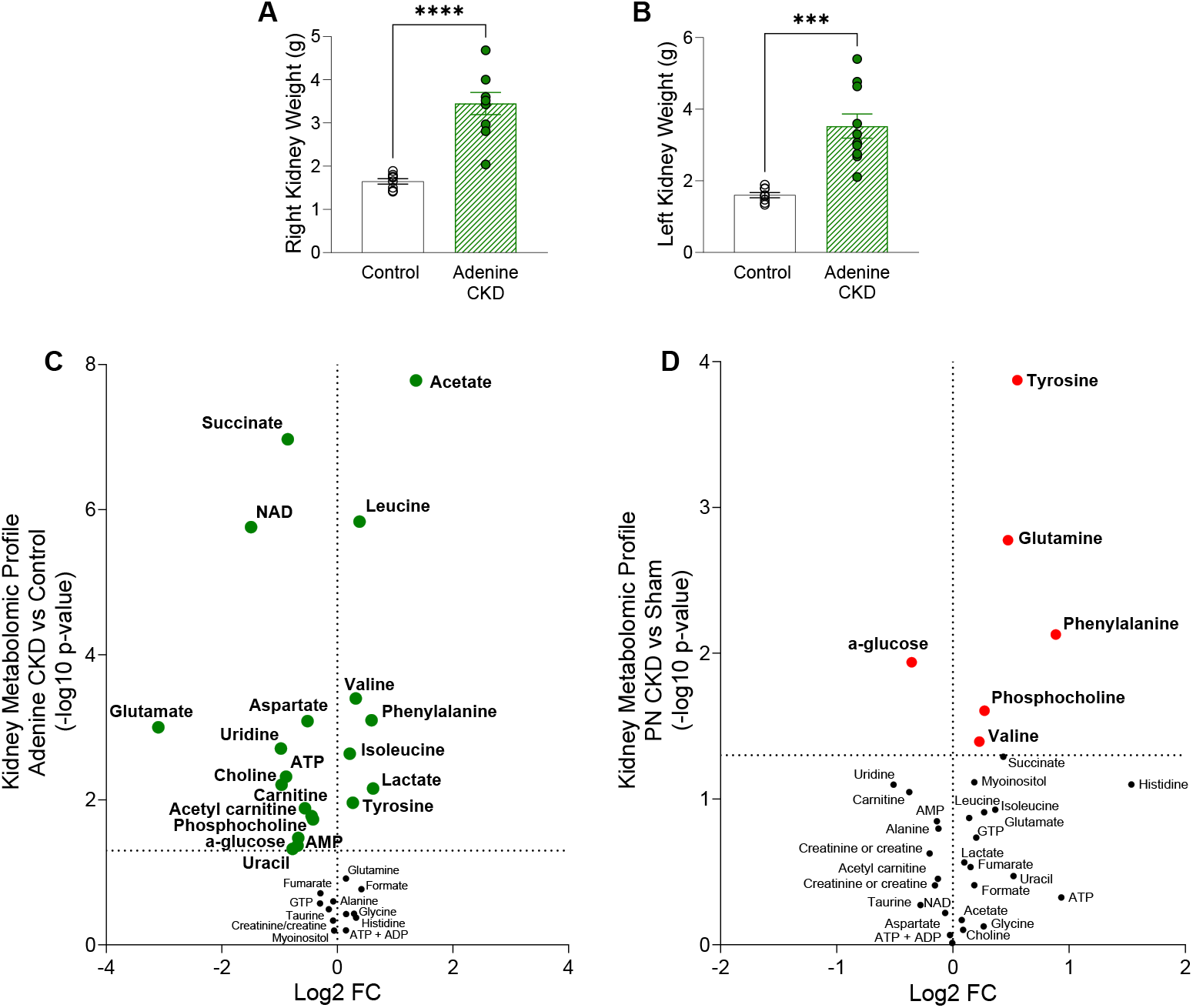
Kidney morphology and metabolomic profile of adenine and PN CKD models. (**A-B)** Right and left kidney weight. Analysed by unequal variance t-test, *p<0.05, control n=8, Adenine n=10. **(C-D)** Kidney metabolomic profile measured by ^1^H NMR spectroscopy of (D) adenine diet CKD (n=9) vs control diet (n=5) and (E) PN CKD (n=4) vs sham (n=4). Each point represents the fold change (FC) of a metabolite plotted against the associated level of statistical significance for the change analysed by t-test (horizontal dashed line indicates p=0.05 threshold). Data displayed as mean ± SEM.

Both experimental models of CKD had marked changes in the metabolomic profile of kidney tissue, to a greater extent in the adenine model (Fig.5C, D). In the adenine CKD model overt metabolic changes were observed with 20 changed out of 32 metabolites studied, including reduction in energetics (reduced ATP, Fig.5C). Energetics were unaffected in the PN CKD model (Fig.5D). There are similarities in the kidney metabolic remodelling between both CKD models including increased amino acids (Fig.5C-D).

#### Liver

The liver phenotype of both CKD models included reduction in liver weight (Fig.6A-B). There were changes in the liver metabolomic profile of adenine CKD animals including a reduction in taurine (Fig.6C). In PN CKD, changes in the metabolomic profile included changes in energy reserve (Fig.6D).

**Fig. 6.**
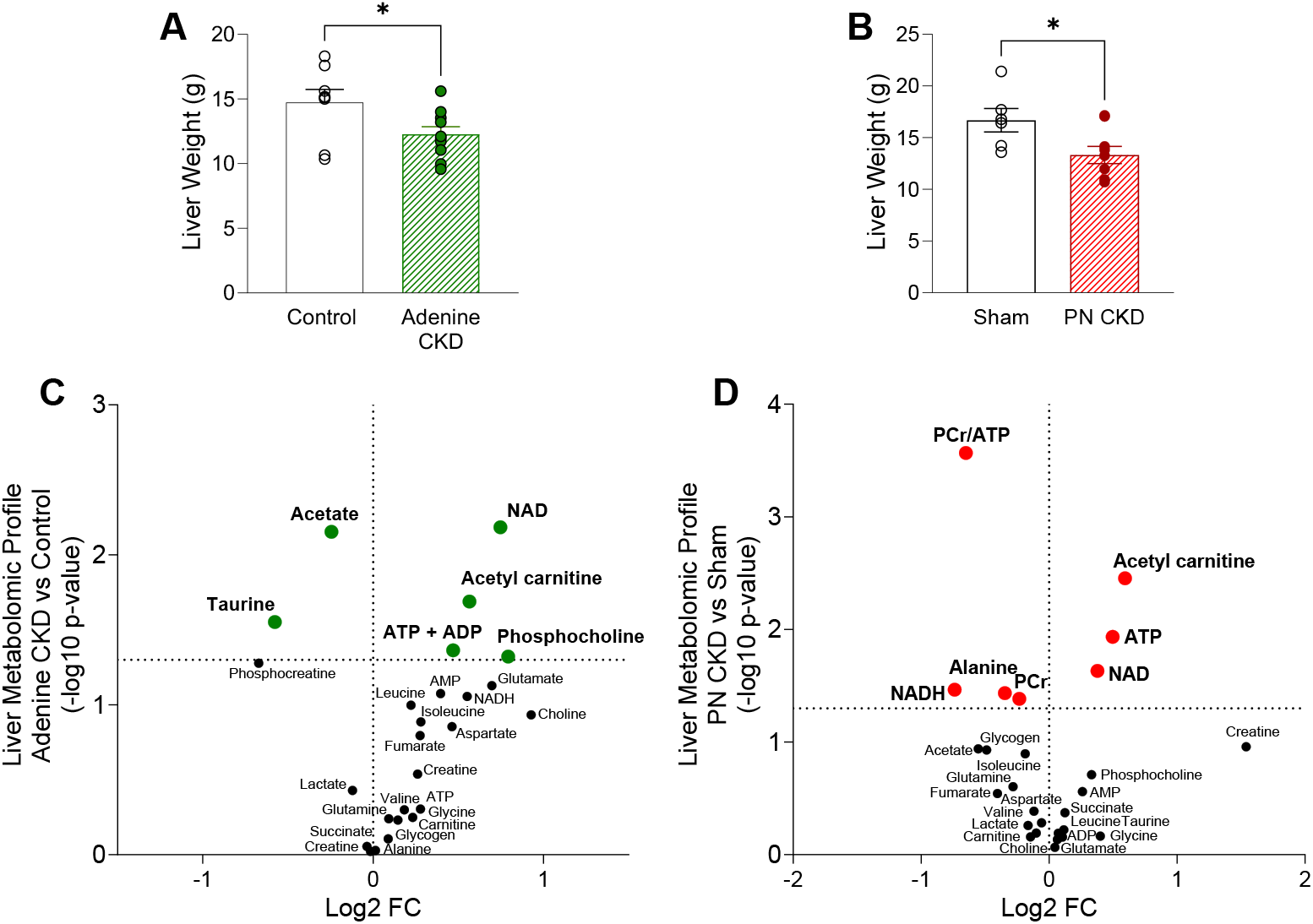
Metabolomic profile of liver in adenine and PN CKD models. (**A-B**) Liver weight in (A) adenine diet CKD (n=10) vs control diet (n=8) animals and (B) PN CKD (n=9) and sham (n=6) animals. Analysed by Student’s t-test, *p<0.05. (**C-D**) Liver metabolomic profile measured by ^1^H NMR spectroscopy of (C) adenine diet CKD (n=5) vs control diet (n=4) and (D) PN CKD (n=4) vs sham (n=4). Each point represents the fold change (FC) of a metabolite plotted against the associated level of statistical significance for the change analysed by t-test (horizontal dashed line indicates p=0.05 threshold). Data displayed as mean ± SEM.

### ^31^P MRS assessment of exercising leg muscle of CKD patients identifies functional bioenergetic impairment

Given the finding of perturbed metabolism in the peripheral tissues of the CKD animal models, we sought to measure the energetic reserve in skeletal muscle of CKD patients when challenged with exercise. The demographic and clinical characteristics of the CKD patients and healthy control study participants is shown in Table 3. ^31^P MRS was obtained from the tibialis anterior muscle at baseline, during exercise, and during the 10-minute recovery period (Fig.7A). The depletion of PCr can be observed across the exercise period, followed by resynthesis during recovery (Fig.7B-C). At baseline, CKD patients had a comparable PCr/ATP ratio to healthy controls (Fig.7D), however the utilisation of PCr during exercise was 35% lower in the CKD patient group (ΔPCr, Fig.7E). Furthermore, we examined the maximum torque generation during exercise and found the CKD patient group generated significantly lower force than healthy controls (Fig.7F). Whilst the muscle PCr utilisation is comparable between CKD patients and controls when torque generation is accounted for (Fig.7G), mitochondrial capacity (Qmax) is significantly decreased (Fig.7H) alongside a decreased initial rate of PCr resynthesis (Fig.7I). The overall PCr recovery indicated by Tau as well as pH was comparable between CKD patients and healthy controls (Fig.S7A-B).

**Fig. 7.**
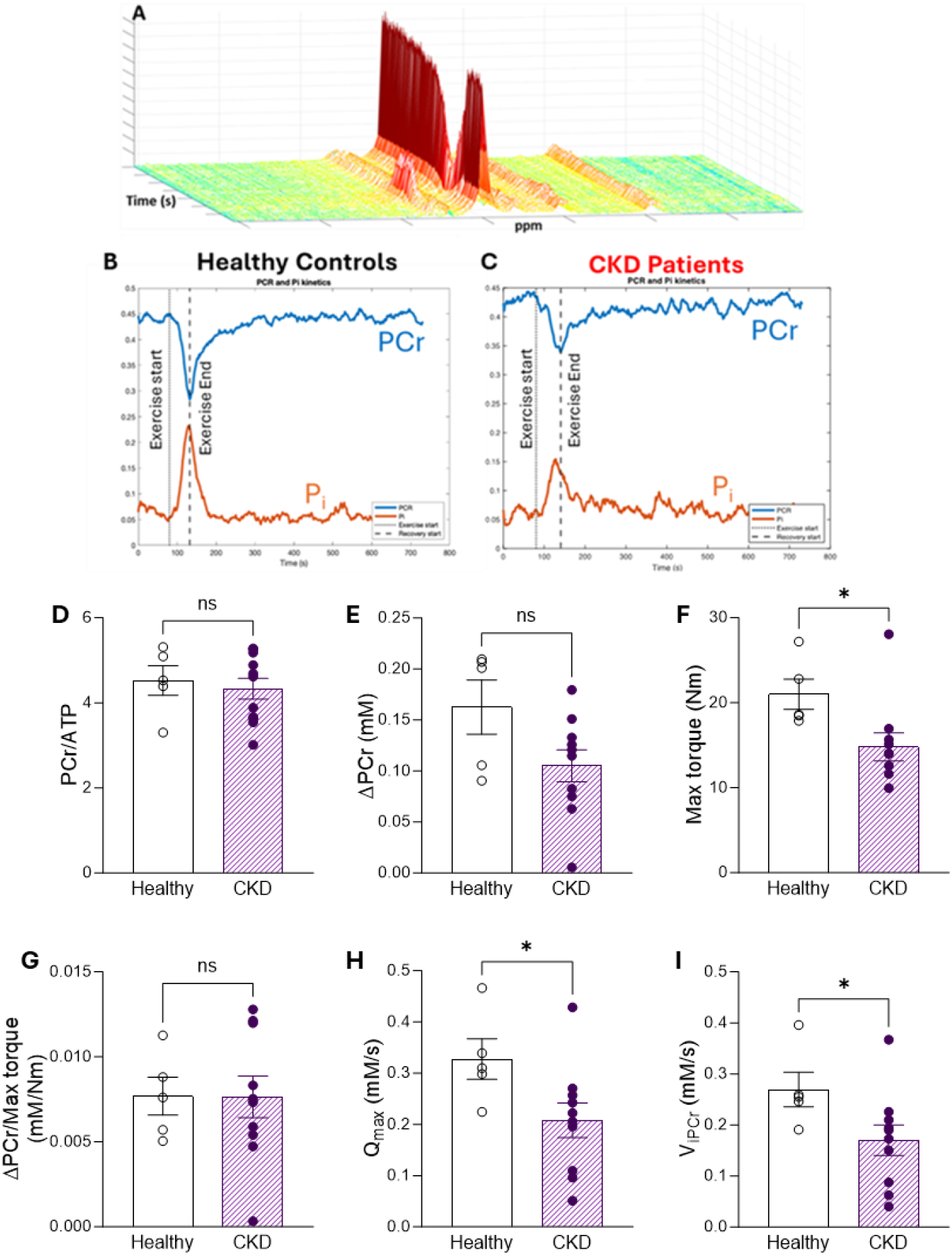
Human ^31^P MRS in tibialis anterior muscle of healthy controls and CKD patients during exercise. (**A**) Dynamic ^31^P spectra. (**B-C**) Change in PCr and Pi before, during and after 30 seconds of exercise. (**D**) Baseline PCr levels, (**E**) ΔPCr (difference in PCr between rest and end of exercise), (**F**) Maximum torque generation during exercise and (**G**) ΔPCr normalised to maximum torque generation in CKD patients and healthy controls. (**H**) Estimated mitochondrial capacity for maximal ATP resynthesis (Q_max_), determined as described in the text. (**I**) Estimated initial rate of PCr resynthesis following exercise as described in the test. (D-I) *p<0.05, healthy controls n=5, CKD patients n=10. (D, G-H) Analysed by unequal variance *t*-test, (E-F, I) analysed by Mann-Whitney U test. Data displayed as mean ± SEM.

**Table 3.**
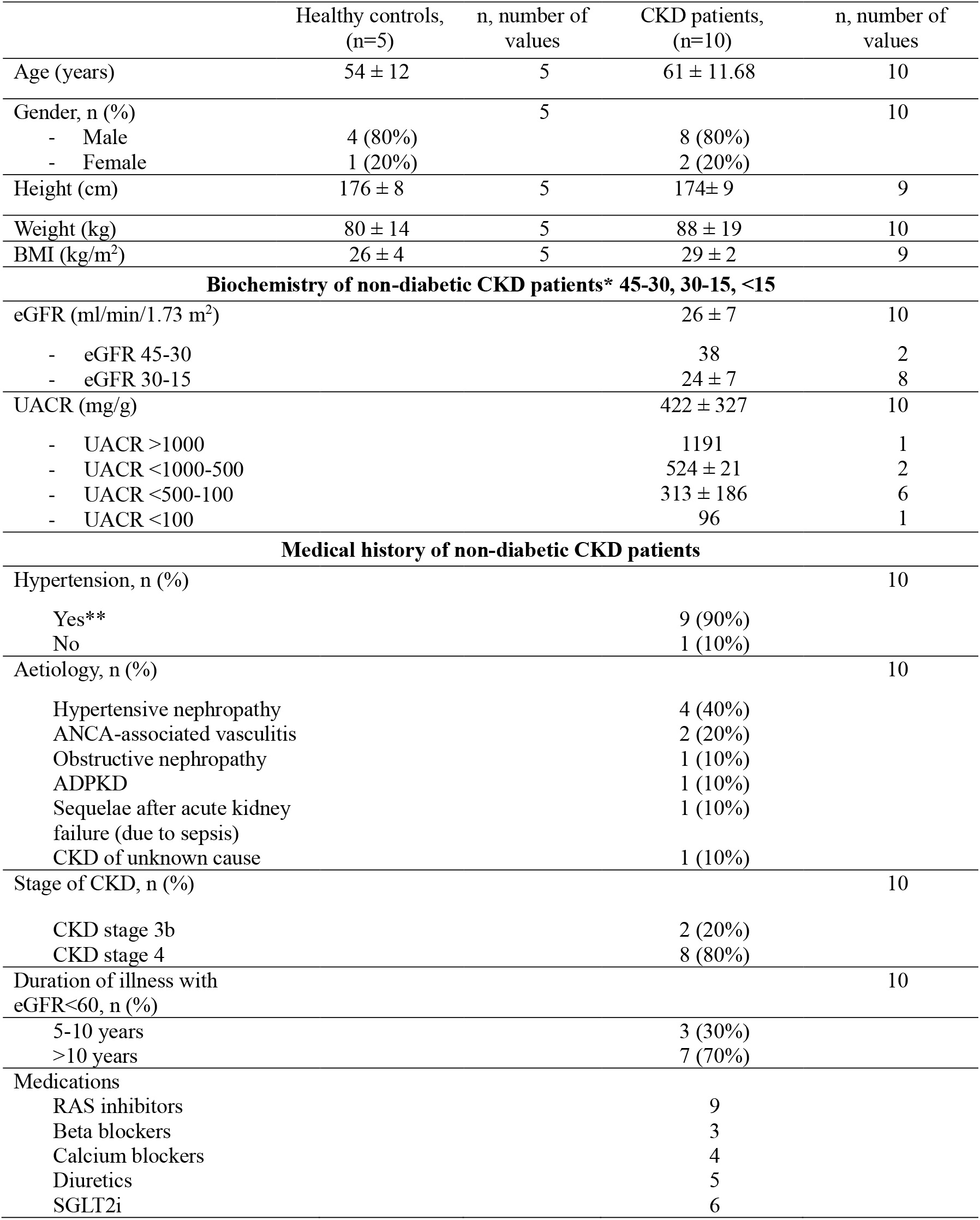
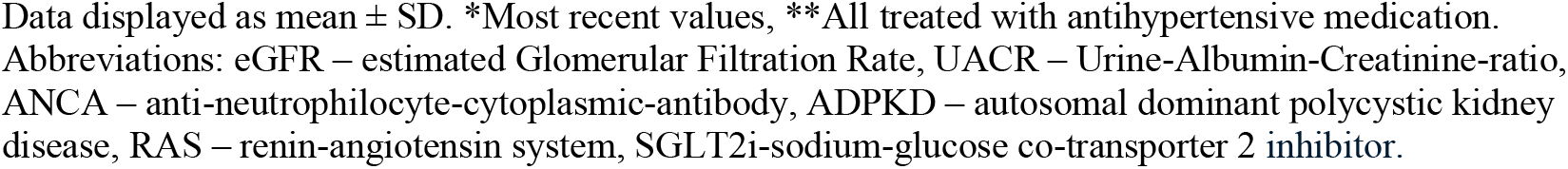
Demographics of CKD patients and healthy controls.

|  | Healthy controls,<br>(n=5) | n, number of<br>values | CKD patients,<br>(n=10) | n, number of<br>values |
| --- | --- | --- | --- | --- |
| Age (years) | 54 ± 12 | 5 | 61 ± 11.68 | 10 |
| Gender, n (%) |  | 5 |  | 10 |
| - Male | 4 (80%) |  | 8 (80%) |  |
| - Female | 1 (20%) |  | 2 (20%) |  |
| Height (cm) | 176 ± 8 | 5 | 174 ± 9 | 9 |
| Weight (kg) | 80 ± 14 | 5 | 88 ± 19 | 10 |
| BMI (kg/m<sup>2</sup>) | 26 ± 4 | 5 | 29 ± 2 | 9 |
| **Biochemistry of non-diabetic CKD patients\* 45-30, 30-15, <15** |  |  |  |  |
| eGFR (ml/min/1.73 m<sup>2</sup>) |  |  | 26 ± 7 | 10 |
| - eGFR 45-30 |  |  | 38 | 2 |
| - eGFR 30-15 |  |  | 24 ± 7 | 8 |
| UACR (mg/g) |  |  | 422 ± 327 | 10 |
| - UACR >1000 |  |  | 1191 | 1 |
| - UACR <1000-500 |  |  | 524 ± 21 | 2 |
| - UACR <500-100 |  |  | 313 ± 186 | 6 |
| - UACR <100 |  |  | 96 | 1 |
| **Medical history of non-diabetic CKD patients** |  |  |  |  |
| Hypertension, n (%) |  |  |  | 10 |
| Yes** |  |  | 9 (90%) |  |
| No |  |  | 1 (10%) |  |
| Aetiology, n (%) |  |  |  | 10 |
| Hypertensive nephropathy |  |  | 4 (40%) |  |
| ANCA-associated vasculitis |  |  | 2 (20%) |  |
| Obstructive nephropathy |  |  | 1 (10%) |  |
| ADPKD |  |  | 1 (10%) |  |
| Sequelae after acute kidney failure (due to sepsis) |  |  | 1 (10%) |  |
| CKD of unknown cause |  |  | 1 (10%) |  |
| Stage of CKD, n (%) |  |  |  | 10 |
| CKD stage 3b |  |  | 2 (20%) |  |
| CKD stage 4 |  |  | 8 (80%) |  |
| Duration of illness with eGFR<60, n (%) |  |  |  | 10 |
| 5-10 years |  |  | 3 (30%) |  |
| >10 years |  |  | 7 (70%) |  |
| Medications |  |  |  |  |
| RAS inhibitors |  |  | 9 |  |
| Beta blockers |  |  | 3 |  |
| Calcium blockers |  |  | 4 |  |
| Diuretics |  |  | 5 |  |
| SGLT2i |  |  | 6 |  |

Data displayed as mean ± SD. \*Most recent values, \*\*All treated with antihypertensive medication. Abbreviations: eGFR – estimated Glomerular Filtration Rate, UACR – Urine-Albumin-Creatinine-ratio, ANCA – anti-neutrophilocyte-cytoplasmic-antibody, ADPKD – autosomal dominant polycystic kidney disease, RAS – renin-angiotensin system, SGLT2i-sodium-glucose co-transporter 2 inhibitor.

## DISCUSSION

Given the systemic nature of CKD and metabolic crosstalk between organs, we investigated both the cardiac and systemic metabolic signature (skeletal muscle, liver, kidney) of two CKD animal models with two distinct aetiologies, durations and severities: tubulointerstitial fibrosis and glomerulosclerosis. Moreover, we assessed the metabolic capacity of skeletal muscle of human CKD patients. This study identified that CKD, regardless of aetiology, leads to cardiometabolic remodelling and increased susceptibility to ischaemia reperfusion injury. However, changes in systemic metabolomic profile (muscle, liver, kidney) exceed severity of cardiac metabolic remodelling. We also show that despite comparable energetic capacity to healthy controls at rest, CKD patients have an inability to utilise their muscle energetic reserve. Therefore, our study shows inability of both heart and skeletal muscles in CKD to respond to a metabolic stress.

Adenine-diet induced CKD is of distinct aetiology to PN and induces CKD with the absence of hypertension [12]. However, cardiometabolic remodelling is less extensively reported for the adenine-diet induced CKD experimental model. The adenine CKD model caused a more severe perturbation of renal function and anaemia, as well as significantly impaired systemic glucose homeostasis (hypoglycaemia, hypoinsulinaemia) not seen in the PN CKD model. The absence of pressure overload is evidenced by the lack of LVH. Observed cardiac functional changes are suggestive of compensatory functional response to maintain ejection fraction under hyperdynamic confounders including severe anaemia.

Decline in kidney function impacts the reabsorption and therefore availability of metabolites within the body. Metabolic profile of adenine-diet induced CKD cardiomyopathy was characterised by reduced exogenous glucose supply due to circulating hypoglycaemia and hypoinsulinaemia which could be responsible for reduced expression of insulin-responsive myocardial glucose transporter GLUT4. Increased expression of fatty acid transporter CD36 accompanied by increased myocardial fatty acid metabolism intermediate (acetyl carnitine) are indicative of increased myocardial uptake and utilisation of fatty acids for the maintenance of cardiac energy reserve as PCr/ATP and glycogen content were maintained. Depleted myocardial lipid pools are also indicative of increased use of fatty acids. Given there is no energetic deficit and no increase in BNP, there is no evidence of HF in adenine diet-induced CKD.

We have previously shown that experimental uraemia induced by PN leads to development of anaemia, hypertension, LVH and substrate utilisation switch from fatty acid to glucose utilisation concurrent with a decline in energy reserve [6, 13-17]. Here we show that although there are no extensive changes in intermediate myocardial metabolite concentration in PN CKD, alterations in myocardial amino acid profile are indicative of metabolic adaptation to maintain ATP supply [18]. However, these metabolic adaptations were insufficient to maintain a healthy cardiac phenotype as reduced PCr/ATP and elevated BNP levels are indicative of progression into HF with a preservation of ejection fraction.

Any underlying cardiac metabolic adaptations in CKD may be sufficient to maintain energetic reserve or cardiac function under basal conditions, but the ability to respond to stress is ultimately impaired. We found that irrespective of the renal failure aetiology and the state of energy reserve of the heart, the presence of CKD with its multifaceted pathophysiology enhances the myocardial susceptibility to ischaemic stress. Adenine CKD hearts show recovery during the initial 5 minutes of reperfusion reflective of the preserved baseline energetics but ultimately recovery during reperfusion is poor. This increased susceptibility to ischaemia reperfusion injury is in line with previously reported enhanced susceptibility to dobutamine stress, increased susceptibility to mitochondrial permeability transition pore opening in CKD models as well as increased incidence of ischaemic cardiac death in CKD patients [6, 19-21].

Muscle wastage, fatigue and poor physical performance are common in CKD, and a major debilitating feature impacting patients’ quality of life [22, 23]. Changes in the adenine model CKD such as reduced body weight and increased circulating creatine kinase are indicative of muscle damage. We also found that a muscle specific decline in PCr/ATP ratio in the adenine diet-induced CKD model preceded decline in cardiac energetic reserve. This suggests that in CKD, skeletal muscle energetic impairment precedes detrimental cardiac changes. Previous work has shown skeletal muscle energetic reserve correlates with cardiac energetic decline in experimental uraemia, highlighting the metabolic and functional relationship between organs [6]. We investigated this relationship further in CKD patients. We found that despite comparable baseline energy reserve, during exercise CKD patients were unable to utilise PCr to the same extent as healthy controls. Furthermore, torque generation was lower in CKD patients versus healthy controls. Given the relationship between CKD, muscular wastage and fatigue this may be reflected in reduced capacity for muscle force generation observed. Our findings are in line with previous reports that CKD patients have reduced peak oxygen capacity, skeletal muscle mitochondrial capacity, reduced mitochondrial number, mitochondrial dysfunction and increased superoxide generation [24-26]. Furthermore, in the calf muscle of haemodialysis patients, ATP supply via oxidative metabolism was shown to be impaired and compensated for by an increase in anaerobic glycolysis [27]. We found that the utilisation of PCr relative to the torque generation was comparable. Therefore, the energetic impairment in skeletal muscle of CKD patients is not solely governed by the availability of PCr nor by a need to utilize greater amounts of energy to produce the same level of force. The limiting factor appears to be an inability to utilise the available energy reserve, leading to the development of fatigue.

Kidney metabolome was perturbed in both experimental models. Severe renal injury caused by adenine deposits in the kidney [12] led to extensive metabolic alterations including reduced kidney energy reserve. Amongst the multiple systemic complications in CKD, our study also identified the interplay between kidney and liver. Reduction in liver weight as well as increase in plasma liver function enzyme marker is indicative of chronic liver damage. Both CKD models are characterised by extensive alterations in liver redox, energy reserve and elevation of the lipid metabolism intermediates. Furthermore, a reduction in taurine was noted in liver from adenine CKD models, a key regulator of hepatic oxidative stress and fatty liver disease [28]. CKD is associated with higher prevalence of liver metabolic diseases such as cirrhosis and non-alcoholic fatty liver disease as two pathologies share several risk factors, including insulin resistance, oxidative stress, inflammation alongside an increased prevalence of cardiovascular disease [10, 29, 30].

How do systemic metabolic confounders common to both models predispose to poor cardiovascular outcomes in CKD? Renal dysfunction is known to lead to accumulation of uraemic toxins such as indoxyl sulfate, kynurenic acid, indole-3-acetic acid, p-cresol sulfate which have been shown to directly inhibit mitochondrial oxidative phosphorylation, cause metabolic perturbation and insulin resistance in peripheral tissues [25, 31, 32] (Fig.8). Moreover, severely decreased haematocrit (anaemia) impairs O_2_ delivery resulting in organ hypoxia [33] further exacerbating metabolic remodelling, mitochondrial dysfunction thus leading to energetic deficit (Fig.8). Loss of glucose through impaired renal reabsorption also leads to reduced plasma availability. Collectively, this perfect ‘storm’ of events associated with CKD impacts systemic metabolism, triggers organ dysfunction and predisposes hearts to remodelling (Figure 8).

**Fig. 8.**
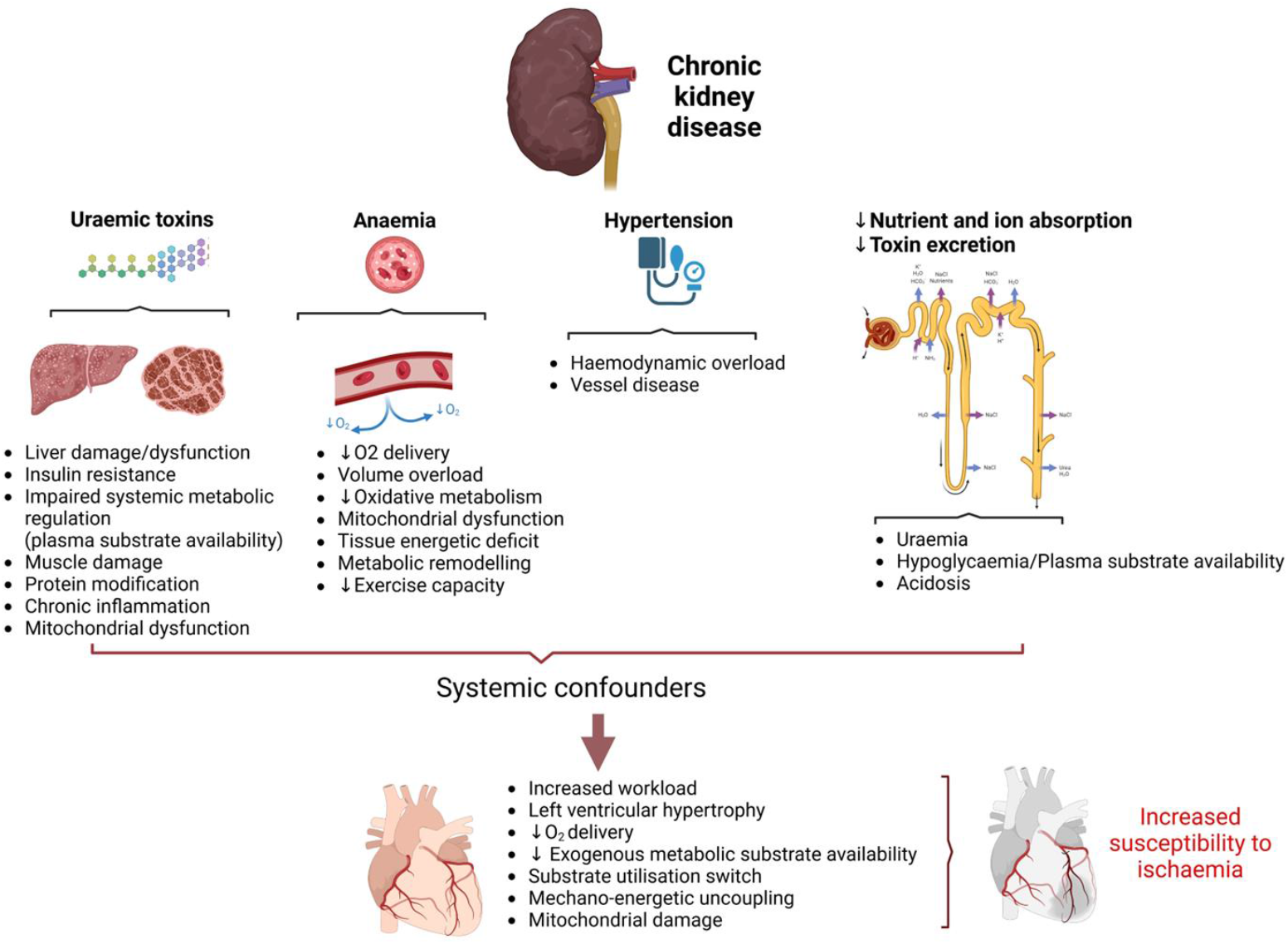
The impact of the systemic confounders on cardiac outcomes in CKD – mechanistic summary

Crosstalk between organs has been identified in HFpEF of different aetiologies [34] and likely contributes to the systemic nature of CKD including the development of uraemic cardiomyopathy. Improved understanding of organ interplay can enhance monitoring protocols and pave the way for interventions that can mitigate the adverse effects of CKD on overall metabolic health. Moreover, therapeutic targeting and prevention of systemic metabolic derangement may be a new approach to ameliorate aberrant cardiac remodelling in CKD.

## MATERIALS AND METHODS

### Animals

250 g male Wistar rats (n=43) were purchased from Charles River Laboratories (UK). All animal experiments were in keeping with the UK Home Office Animals (Scientific Procedures), 1986.

### Experimental models of chronic kidney disease

Partial nephrectomy (PN) CKD was induced using an established two-stage surgical procedure for a period of 12 weeks [12, 15, 16]. Adenine diet model of CKD was induced by 4 week adenine dietary protocol (0.75% adenine in standard rat chow) followed by 4 week standard chow diet [35].

### Transthoracic echocardiography

In vivo cardiac function was assessed by echocardiography (Vevo 770;Visualsonics, Supplementary Materials).

### Biochemical analysis

Tissues (liver, skeletal muscle, heart, kidney) were rapidly collected upon termination and snap frozen in liquid N_2_ [36]. Metabolomic profile was analysed using ^1^H NMR spectroscopy [36, 37]. Plasma obtained from thoracic cavity blood samples taken at the time of sacrifice were analysed by the Clinical Biochemistry Laboratory (Addenbrookes Hospital, Cambridge).

### qPCR

RNA was isolated (Cat:74704;Qiagen) and reverse transcribed (Cat:4374967;Applied Biosystems). qPCR reactions were run in triplicates using SYBR green (Cat:600892;Agilent) on QuantStudio 5 (Applied Biosystems). Gene expression displayed relative to 36B4. Full methods and primer sequences in Supplementary Materials.

### Langendorff heart perfusions

Hearts were rapidly excised, cannulated and perfused in Langendorff mode as previously described [18, 38]. After 20 minutes of equilibration, hearts were subject to 25 minutes global normothermic ischaemia (37°C) and 25 minutes reperfusion.

### Assessment of the human tibialis anterior muscle bioenergetic reserve by dynamic ^31^P-MRS

The clinical study protocol was approved by the Ethics Committee of Central Denmark (ref. 1-10-72-145-23). All subjects provided written consent prior to participation. Ten patients with non-diabetic CKD stage 3b and 4 and five age-matched healthy volunteers with normal renal function (Exclusion criteria in Supplementary Material) were recruited for ^31^P dynamic MRS scanning of the tibialis anterior muscle during exercise [39]. Scans were acquired at a 2 min resting baseline prior to 30 seconds of exercise followed by 10 minutes recovery. Full methods can be found in Supplementary Materials.

### Statistics

Statistical analysis was performed using GraphPad Prism software (version 10.1.0). Normality of data distribution was examined. Comparison between two groups was performed by Student’s t-test, with Welch’s correction for unequal variance applied accordingly, or Mann-Whitney U test used when data were non-normally distributed. A two-way ANOVA was used when two factors were present with post-hoc analysis completed if significant interaction present. Significance accepted when p<0.05.

## Data availability

Data available on reasonable request from the corresponding author.

## Author contributions

Conceptualisation: DA, MMY, NHB, CL

Methodology: DA, MY, MA, NP, FC, ESSH, JEC, TRE, MV, LK, LV, JJJJM, NHB, CL

Investigation: DA, MY, MA, NP, FC, ESSH, JEC, TRE, MV, LK, LV, JJJJM, NHB

Visualisation: MY, DA, MA

Funding acquisition: DA, MMY, CL,

NHB Project administration: DA

Supervision: DA, MMY, JJJJM, NHB, CL

Writing – original draft: MY, MA, DA

Writing – review & editing: MY, DA, MMY, JJJJM, MA, ESSH, TRE, MV, LV, NHB, CL

## Acknowledgments

We would like to thank Dr Nasima Kanwal and School of Physical and Chemical Sciences NMR Facility, Queen Mary University of London. DA is a co-PI of Barts Metabolism Network. We acknowledge the following sources of funding: Wellcome Trust Career Re-Entry Fellowship 221604/Z/20/Z (DA), Barts Charity grant G-002145 (DA), British Heart Foundation MRes Studentship FS/4YPhD/P/20/34016 (NP), Barts HCA Fellowship (LK), Leventis Foundation Cyprus (LK), Novo Fonden grant NNF21OC0068683 (MA, JJM, CL, MV), MRC MDU Mouse Biochemistry Laboratory MC_UU_00014/5, Centre of Excellence in Medical Engineering funded by Wellcome Trust (TRE), EPSRC WT/203148/Z/16/Z (TRE), BHF Centre of Research Excellence RE/24/130035 (TRE), Wellcome Trust Sir Henry Dale Fellowship 221805/Z/20/Z (LV), Slovak grant agencies VEGA 2/0004/23 and APVV 21-0299 (LV).

## Notes

### Competing Interest Statement

The authors have declared no competing interest.

### Summary of Updates

We have updated manuscript abstract and discussion

