## Supplemental Material for "Altered systemic bioenergetic reserve in chronic kidney disease predisposes hearts to worse functional outcomes"

Supplementary material

**Transthoracic echocardiography**

For the assessment of cardiac function, echocardiography was performed in control diet, sham control, adenine diet induced-CKD and PN CKD animals. Anaesthesia was induced with 5% isoflurane and maintained at 1.5-2% for the duration of the procedure. Before assessment of cardiac function, fur was removed from the chest area to allow accurate assessment of cardiac function and rats were allowed to stabilise prior to recording. Body temperature was maintained at 37 °C. Echocardiography images were recorded using a Vevo 770 imaging system (Visualsonics, Canada). Percentage ejection fraction (EF) was calculated from the M-mode measurements in the parasternal short axis view at the level of the papillary muscles.

**qPCR**

RNA was isolated using RNeasy Fibrous Tissue Kit (Qiagen) according to the manufacturer's instructions. RNA quantity and quality was assessed using NanoDrop ND-1000 spectrophotometer and only RNA with 260/280 >1.8 was used for analysis. 1µg total RNA was reverse transcribed using High-Capacity cDNA Reverse Transcription Kit (Applied Biosystems). For qPCR, gene specific primer sequences are shown in Supplementary Table 1. qPCR reactions were performed in triplicates using Brilliant III Ultra-fast SYBR Green qPCR Master Mix with low ROX (Agilent) according to the manufacturer's instructions with the following cycling conditions: 95°C for 3 min, 40 cycles of 95°C for 5 sec and 60°C for 60 sec using QuantStudio 5 (Applied Biosystems). The relative quantity of each gene was calculated using ΔΔCT method with 36B4 as endogenous control.

**Human Subjects**

Ten patients (8 males, 2 females) with non-diabetic CKD stage 3b and 4, a mean age of 61 years and a mean estimated glomerular filtration rate of 27 ml/min/1.73m$^2$ were recruited for the $^{31}$P dynamic MRS study. Exclusion criteria for all human participants were: diabetes, uncontrolled heart or lung disease, pronounced peripheral vascular disease, severe osteoarthritis in the joints of the lower extremities, neurological and rheumatological disease affecting the muscles, or other disease that reduces the ability to perform physical activity, severe allergies, pregnancy, severe claustrophobia, a pacemaker or metal in the body that is not MR-safe or that interferes with the images, circumference incl. arms >160 cm. Further exclusion criteria for CKD patients were presence of heart failure or ischaemic heart disease, previous clinical cardiovascular events (AMI, stroke incl. TCI case). The clinical characteristics of the patient cohort can be found in Table 3. The control arm consisted of 5 healthy volunteers (4 males, 1 female) with a mean age of 54 years and normal renal function.

**Assessment of the human tibialis anterior muscle bioenergetic reserve by dynamic $^{31}$P-MRS**

Human subjects underwent $^{31}$P dynamic MRS scanning of the lower limb, specifically the tibialis anterior muscle, during exercise [1]. Scans were acquired at a 2 min resting baseline prior to 30 seconds of exercise followed by 10 minutes recovery. Dynamometer exercise in the tibialis anterior muscle was performed using a custom- built MR compatible ergometer. $^{31}$P dynamic spectra (Supplementary Figure 1) were acquired on a 3T MRI scanner (MR750, GE HealthCare, Waukesha, WI, USA) using a $^{31}$P/$^{1}$H dual tuned transmit-receive surface coil (RAPID Biomedical GmbH, Rimpar, Germany). An unlocalised MRS sequence was used to acquire the dynamic spectra; repetition time (TR) = 2000 ms, flip angle (FA) = 70º, repetitions = 360, total acquisition time of 12 min. Data was acquired with 1024 samples at a bandwidth of 5 kHz, corresponding to a spectral resolution of 5 Hz (0.09 ppm). Phosphorus metabolites were determined with Advanced Method for Accurate, Robust, and Efficient Spectral (AMARES) fitting using OXSA [2], an open-source magnetic resonance spectroscopy analysis toolbox in MATLAB. An example of AMARES fitted spectra from the acquired temporal spectra is shown in Supplementary Figure 1. Phosphorus spectra were analysed in MATLAB R2020a (MathWorks, Natick, MA, USA) using the MNS Research Pack created by GE HealthCare. Aerobic muscle load was determined with a pH decrease below 0.5 during exercise using the chemical shift difference of inorganic phosphorus (Pi) and PCr, as previously described [3]: $\mathrm{pH} = 6.75 + \log\left(\frac{\delta - 3.27}{5.69 - \delta}\right)$ where $\delta$ is the ppm difference between PCr and inorganic phosphate as determined through AMARES. Following the recommendations of the $^{31}$P MRS in Skeletal Muscle Expert's Consensus [4], ATP was used as an internal concentration standard assumed constant at 8.2 mM; and the time constant of the phosphocreatine (PCr) recovery following a mono-exponential fit was determined using a least squares minimisation algorithm to the expression

$$\mathrm{PCr(t)} = \mathrm{PCr}_0 + C\left(1 - e^{\frac{-t}{\tau}}\right) \quad [1]$$

where $PCr_0$ is the PCr level at the end of recovery, C is the difference between exercise and recovery end (sometimes called Δ[PCr]), and $\tau$ is the time constant of PCr resynthesis in seconds, a measure of oxidative capacity. From these data, the estimated initial rate of PCr resynthesis was defined as $V_{\mathrm{iPCr}} = \Delta\mathrm{PCr}/\tau$, and $Q_{\mathrm{max}}$, the maximal rate of oxidative ATP synthesis from the PCR recovery data, i.e. a measure of mitochondrial capacity, was determined by a model taking into account the Michaelis-Menten dependence of oxidative metabolism on the concentration of ADP [5]; through the relation

$$[\mathrm{ADP}] = \frac{[\mathrm{Cr}][\mathrm{ATP}]}{[\mathrm{PCr}][\mathrm{H}^+]K_{\mathrm{CK}}} \quad [2]$$

where this relation is solved subject to the assumption of a constant concentration of total creatine, $[\mathrm{tCr}] \approx 42$ mM and utilising knowledge of the Michaelis-Menten coefficient $K_{\mathrm{CK}} \approx 1.66 \times 10^9\ \mathrm{M}^{-1}$.[5] Finally, $Q_{\mathrm{max}}$ was estimated assuming a constant Michaelis-Menten $K_M \approx 30$ µM for the oxidation of ADP, empirically valid in skeletal muscle [6] according to the relation

$$Q_{\mathrm{max}} = V_{\mathrm{iPCr}}\left(1 + \frac{K_M}{[\mathrm{ADP}]_{\mathrm{End\ of\ exercise}}}\right) \quad [3]$$

**Table S1. Primer sequences**

| Primer | Forward Primer Sequence | Reverse Primer Sequence |
|---|---|---|
| 36B4 | AGAGGTGCTGGACATCACAG | CATTGCGGACACCCTCTAG |
| BNP | TGATTCTGCTCCTGCTTTTC | GTGGATTGTTCTGGAGACTG |
| Glut 4 | AGAGGAGAGGGCGGACATT | ACTCTTCATTCAGGCCCTTGG |
| CD36 | GCTGATTACTTCTGTGTAGTAGCTT | TCCTCGTGCAGCAGAATCAA |

**Table S2. Echocardiography parameters for adenine CKD and control diet animals.**

| | Control | Adenine CKD |
|---|---|---|
| Fractional shortening (%) | 48.47 ± 1.76 | 47.67 ± 0.55 |
| Left ventricular internal diameter, end-systole (mm) | 4.13 ± 0.17 | 3.77 ± 0.10 |
| Left ventricular posterior wall, end-systole (mm) | 3.31 ± 0.14 | 3.38 ± 0.14 |
| Left ventricular posterior wall, end-diastole (mm) | 2.21 ± 0.07 | 2.38 ± 0.11 |
| Intraventricular septal thickness, end-systole (mm) | 3.44 ± 0.09 | 3.44 ± 0.09 |
| Intraventricular septal thickness, end-diastole (mm) | 2.14 ± 0.08 | 2.03 ± 0.08 |
| End systolic volume (µl) | 60.45 ± 6.11 | **36.32 ± 1.44*** |

Analysed by Student's t-test, *p<0.05 Adenine vs. control.

**Table S3. Echocardiography parameters for PN CKD and sham animals.**

| | Sham (n=8) | PN CKD (n=7) |
|---|---|---|
| Fractional shortening (%) | 43.32 ± 1.33 | **49.02 ± 1.92*** |
| Left ventricular internal diameter, end-diastole (mm) | 7.81 ± 0.11 | 7.80 ± 0.17 |
| Left ventricular posterior wall, end-systole (mm) | 3.51 ± 0.16 | 3.80 ± 0.09 |
| Left ventricular posterior wall, end-diastole (mm) | 2.39 ± 0.09 | 2.44 ± 0.18 |
| Intraventricular septal thickness, end-diastole (mm) | 2.14 ± 0.06 | 2.35 ± 0.08 |
| End systolic volume (µl) | 89.23 ± 5.06 | **71.03 ± 6.58*** |
| End diastolic volume (µl) | 326.7 ± 17.96 | 320.1 ± 14.95 |

Analysed by Student's t-test, *p<0.05 PN CKD vs. sham.

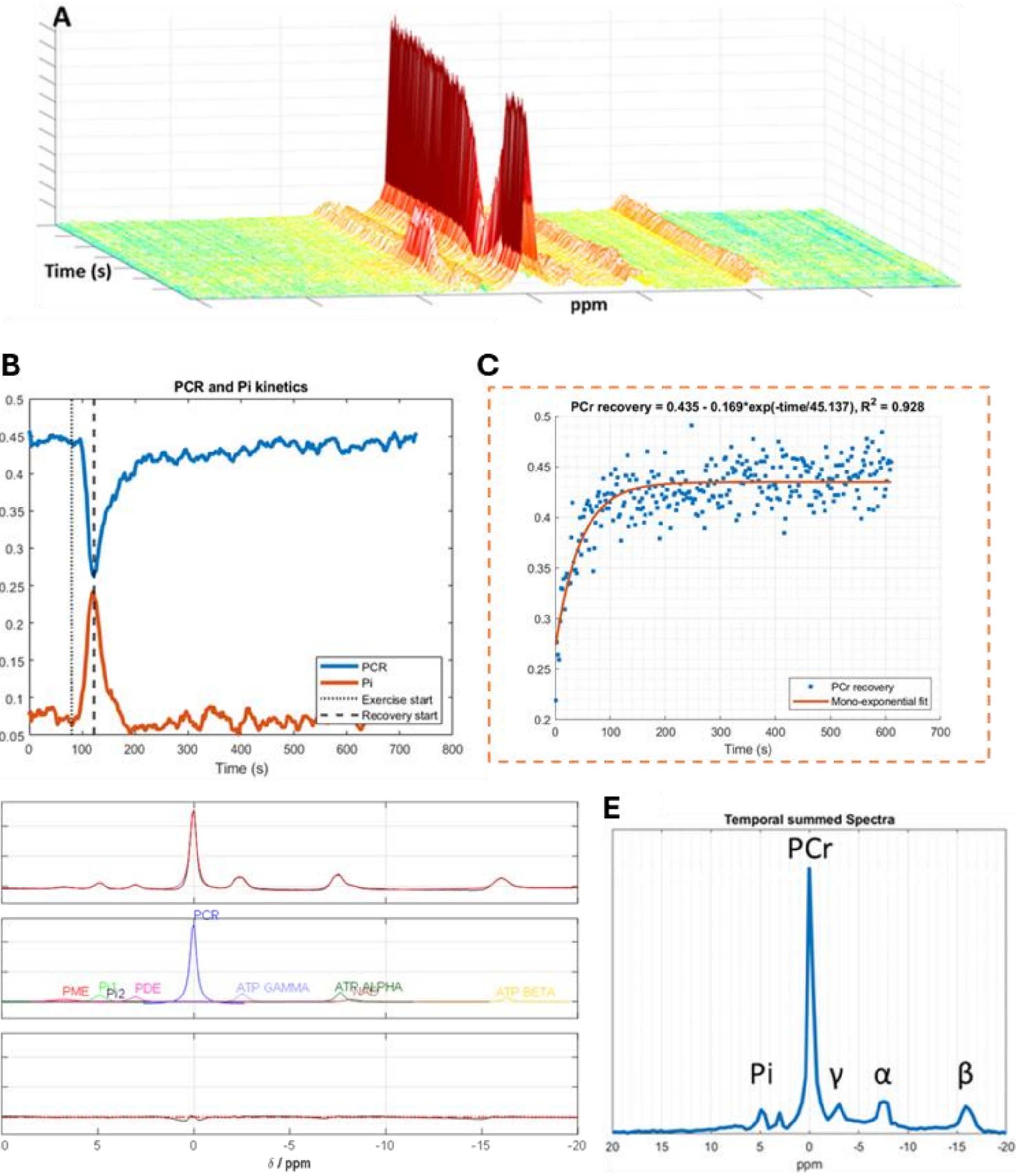


**Fig. S1: Assessment of mitochondrial oxidative capacity by dynamic $^{31}$P-MRS in human tibialis anterior muscle during exercise and recovery.** (**A**) Dynamic $^{31}$P spectra and (**B**) PCr/Pi concentration change in human skeletal muscle at 3T during 2 min baseline, 30 sec exercise, and 10 min recovery with a temporal resolution of 2 s. (**C**) Assessment of oxidative capacity by the time constant of PCr recovery following a mono-exponential fit. (**D**) AMARES fitted temporal summed spectra of the dynamic acquisitions. The plot shows the fitted spectra (top), individual peaks (middle), and fitting residual (bottom). AMARES = Advanced Method for Accurate, Robust, and Efficient Spectral. (**E**) Temporal summed spectra showing metabolites of interest. PCr, phosphocreatine. Pi, inorganic phosphate. ATP alpha (α), beta (β), and gamma (γ).

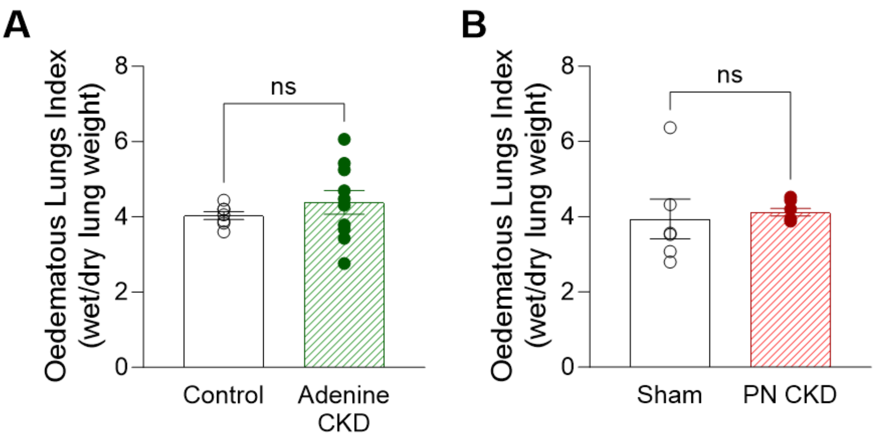


**Fig. S2: Oedematous lung index in CKD models**. Wet lung:dry lung ratio in (**A**) adenine CKD (n=10) vs control (n=7) animals (analysed by unequal variance t-test) and (**B**) PN CKD (n=7) vs sham (n=6) animals (analysed by Mann-Whitney U test).

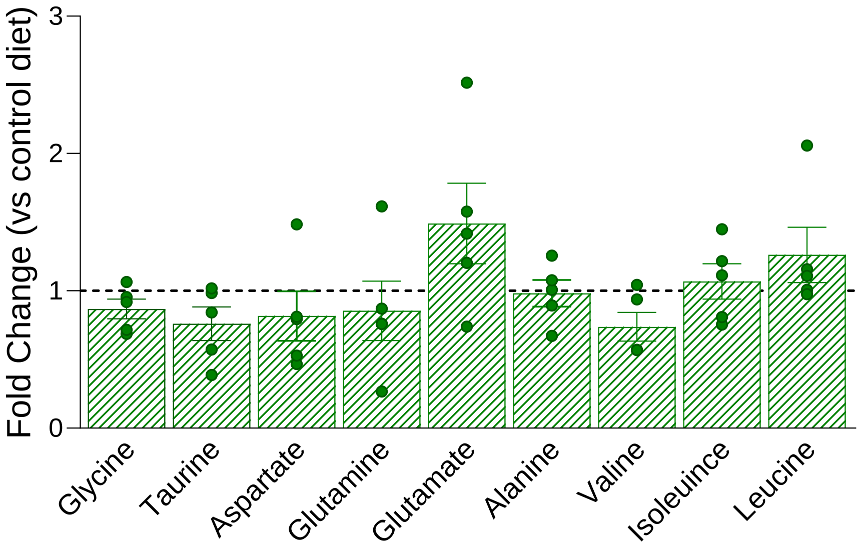


**Fig. S3: Amino acid metabolites in Adenine CKD model.** $^1$H NMR spectroscopy of cardiac tissue from adenine CKD (n=5) expressed as fold change vs control diet (n=9) group. Adenine CKD vs control compared for each metabolite by Student's t-test.

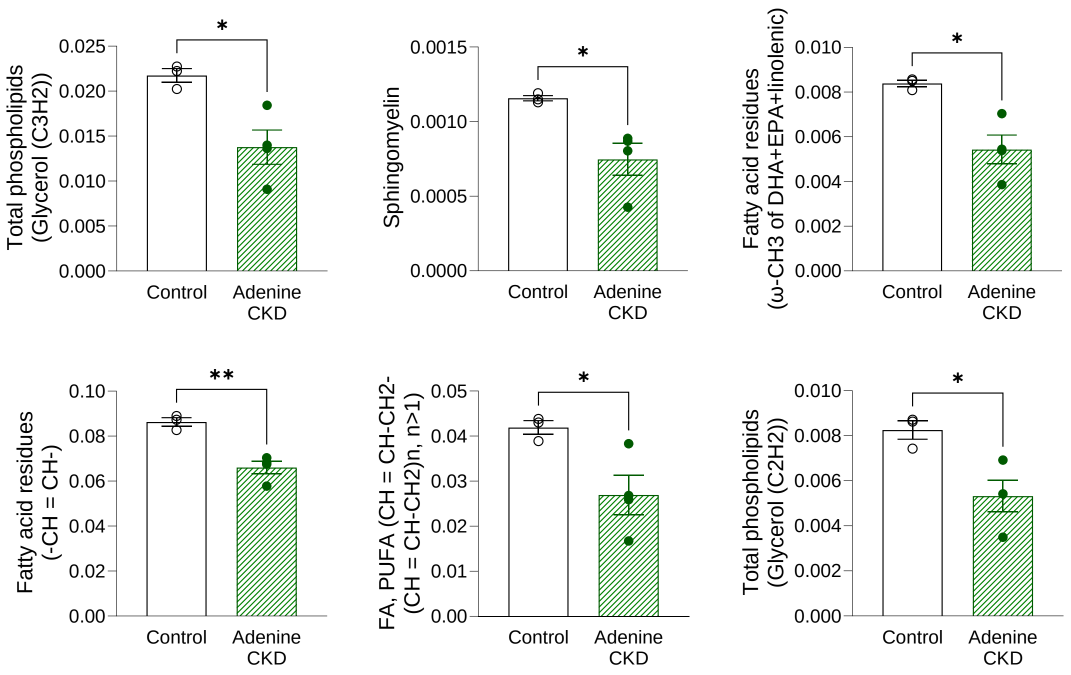


**Fig. S4: Cardiac lipids in adenine induced CKD model.** Cardiac lipids measured by [1]H NMR spectroscopy, Adenine CKD (n=4), control (n=3), *p<0.05 analysed by Student's t-test.

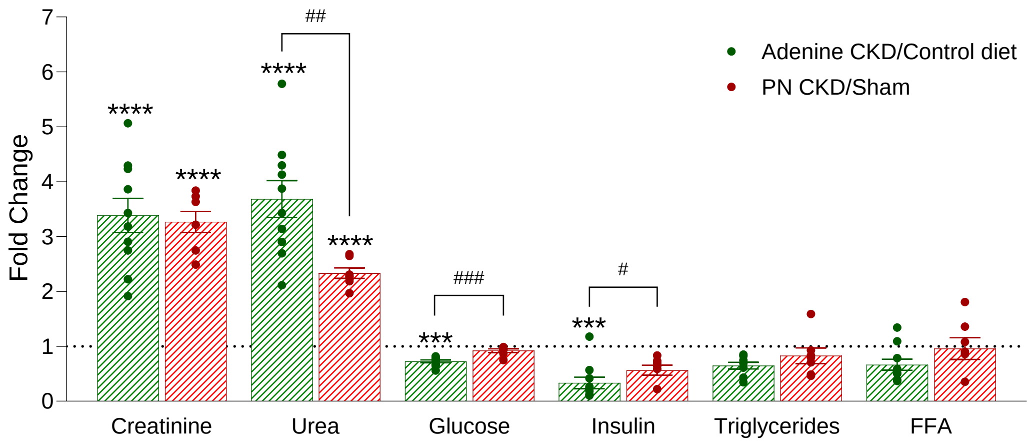


**Fig. S5: Comparison of plasma markers in adenine and PN CKD models**. Plasma levels of Creatinine, Urea, Glucose, Insulin, Triglycerides, and Free fatty acids (FFA) in adenine CKD and PN CKD models (Adenine CKD normalised to control diet and PN CKD normalised to sham control). Difference in CKD model vs respective control displayed above each bar (*p<0.05), analysed by Student's t-test. Difference between normalised adenine CKD and PN CKD displayed with connecting lines (#p<0.05), analysed by Student's t-test.

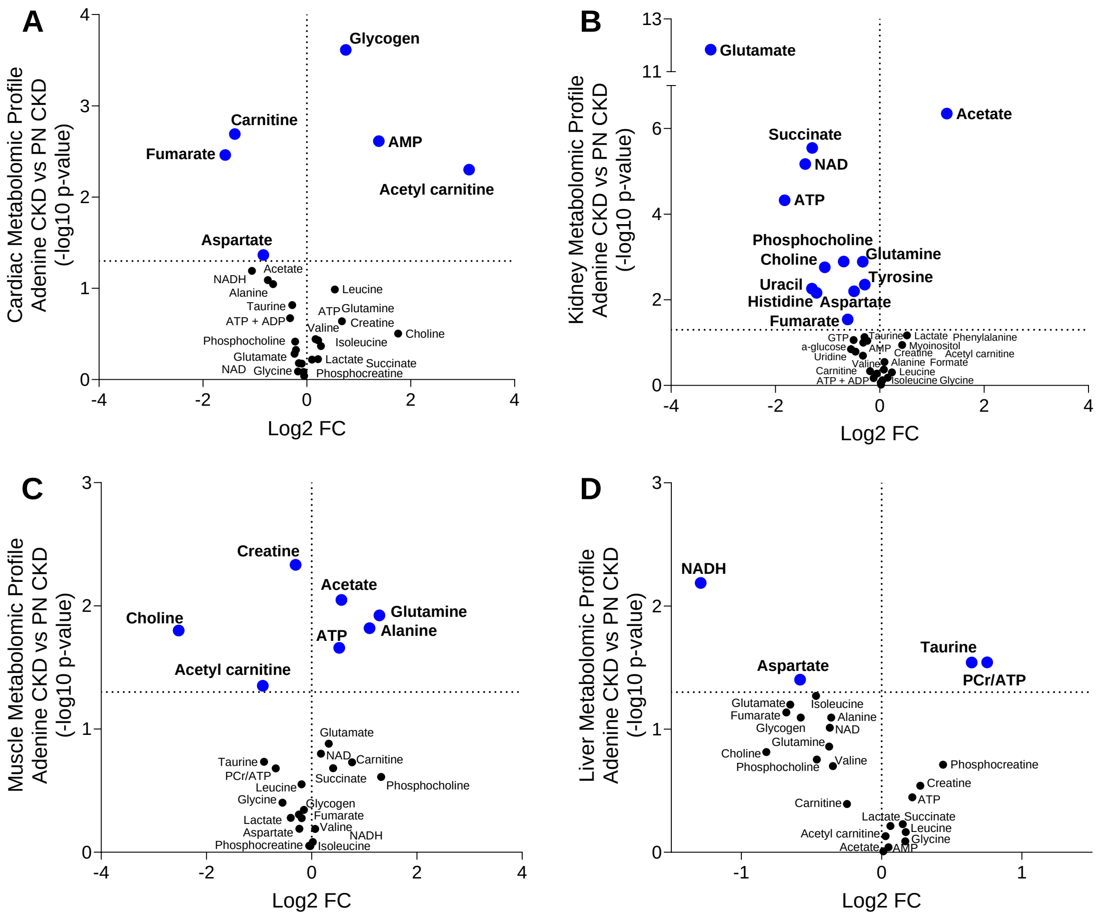


**Fig. S6: Comparison of metabolomic profiles in Adenine diet and PN CKD models.** Metabolomic profile of (**A**) Heart, (**B**) Kidney, (**C**) Muscle and (**D**) Liver in adenine CKD compared to PN CKD, measured by $^{1}$H NMR spectroscopy. Each point represents the fold change (FC) of a metabolite plotted against the associated level of statistical significance for the change analysed by t-test (horizontal dashed line indicates p=0.05 threshold).

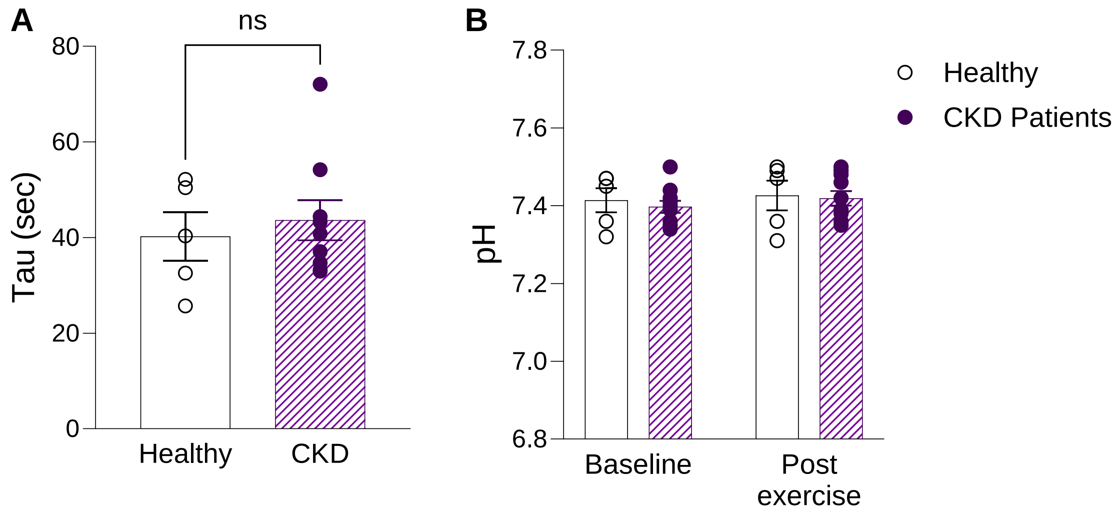


**Fig. S7: Human ³¹P NMR spectroscopy in tibialis anterior muscle in CKD patients during exercise.** (**A**) PCr recovery rate constant following exercise, analysed by Mann Whitney U test. (**B**) pH, measured as the distance between Pi and PCr, before and after 30 seconds of exercise in healthy controls and CKD patients. No significant interaction or main effect, analysed by two-way repeated measures ANOVA. (A-B) Healthy controls n=5, CKD patients n=10.
